# The regulation of a pigmentation gene in the formation of complex color patterns in *Drosophila* abdomens

**DOI:** 10.1101/2020.04.09.034900

**Authors:** Komal K. B. Raja, Mujeeb O. Shittu, Peter M. E. Nouhan, Tessa. E. Steenwinkel, Evan A. Bachman, Prajakta P. Kokate, Alexander H. McQueeney, Elizabeth A. Mundell, Alexandri A. Armentrout, Amber M. Nugent, Thomas Werner

## Abstract

Changes in *cis*-regulatory modules (CRMs) that control developmental gene expression patterns have been implicated in the evolution of animal morphology^1-6^. However, the genetic mechanisms underlying complex morphological traits remain largely unknown. Here we investigated the molecular mechanisms that induce the pigmentation gene *yellow* (*y*) in a complex spot and shade pattern on the abdomen of the quinaria group species *Drosophila guttifera*. We show that the *y* expression pattern is controlled by only one CRM, which contains a stripe-inducing CRM at its core. We identified several developmental genes that may collectively interact with the CRM to orchestrate the patterning in the pupal abdomen of *D. guttifera*. We further show that the core CRM is conserved among *D. guttifera* and the closely related quinaria group species *Drosophila deflecta*, which displays a similarly spotted abdominal pigment pattern. Our data suggest that besides direct activation of patterns in distinct spots, abdominal spot patterns in *Drosophila* species may have evolved through partial repression of an ancestral stripe pattern, leaving isolated spots behind. Abdominal pigment patterns of extant quinaria group species support the partial repression hypothesis and further emphasize the modularity of the *D. guttifera* pattern.

---

How complex morphological features develop and evolve is a question of foremost importance in biology. To address this question, we identified genes underlying abdominal pigmentation pattern development in *Drosophila guttifera* (*D. guttifera*). The abdomen is decorated with six rows of black spots that run along the anterior-posterior axis, divided by a dark dorsal midline shade. This color pattern shows four sub-patterns: a dorsal, median, and lateral pair of spot rows, plus the dorsal midline shade (Fig. 1a, b). *D. guttifera* belongs to the quinaria species group, whose members display highly diverse abdominal pigmentation patterns^7,8^. While *D. guttifera* shows the most complex pattern of this group, most other quinaria group species lack at least one of the four sub-patterns, illustrating the pattern modularity among species. Interestingly, the stripe patterns of certain species often separate into spots^7,8^. In this study, we show that the abdominal pigment patterns of quinaria group members may be formed by a combination of localized spot induction and partial stripe repression.

**Figure 1:**
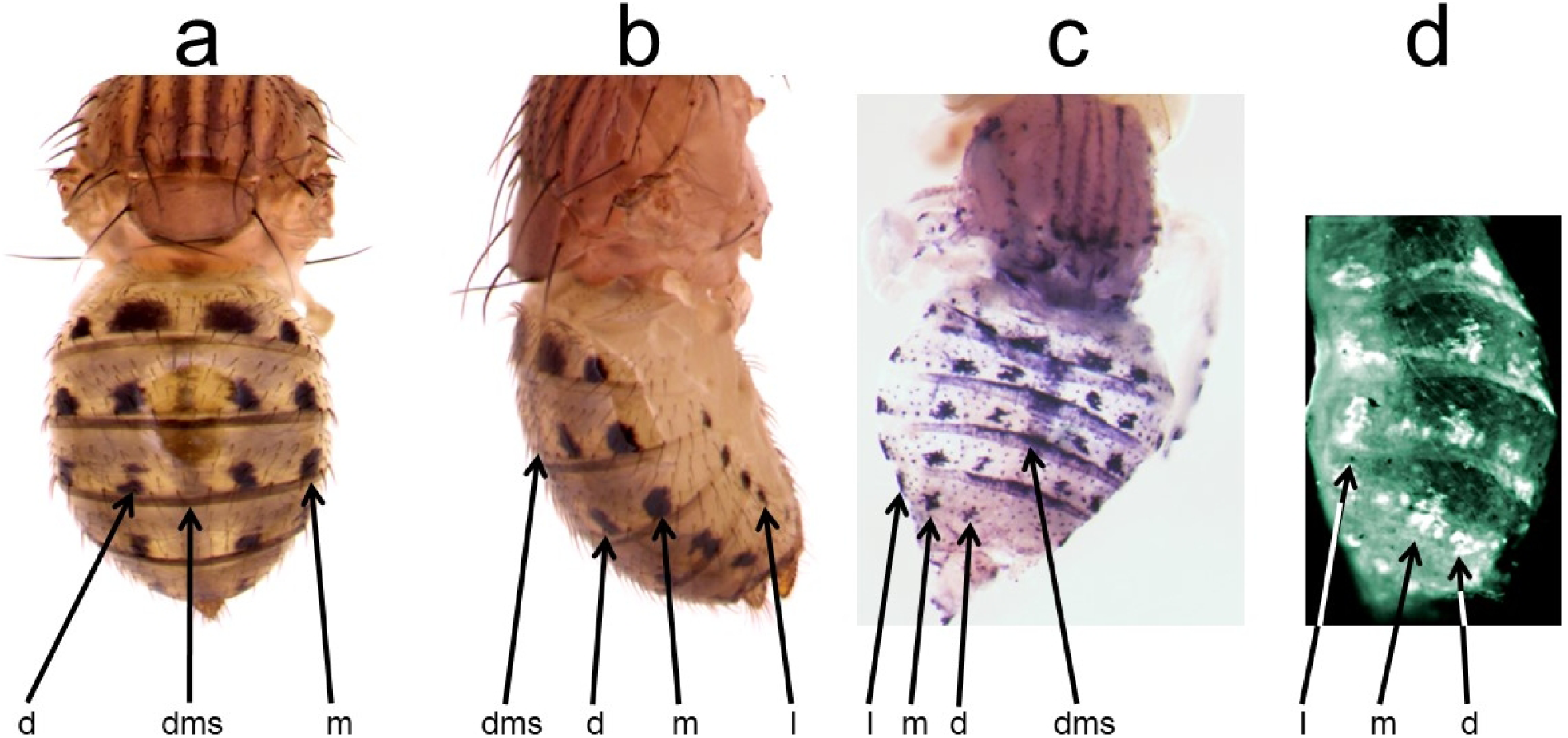
The *D. guttifera* abdominal color pattern is modular. a, Adult, dorsal view. b, Adult, lateral view. c, *yellow* mRNA expression pattern in a pupal abdomen. d, Yellow protein expression pattern in a pupal abdomen. dms = dorsal midline shade, d = dorsal, m = median, l = lateral spot rows.

We focused on the regulation of the *yellow* (*y*) gene, which is required for the formation of black melanin in insects^8-14^. Several *y* gene CRMs have been identified in various *Drosophila* species, and changes in these CRMs and/or in the deployment of *trans*-factors that regulate *y* gene expression have been implicated in the diversification of wing and body pigment patterns^12,15-19^. In *D. guttifera* pupae, *y* gene expression and the location of the Y protein accurately prefigured the complex adult abdominal pigment pattern (Fig. 1c, d). In order to identify putative upstream activators of *y*, we performed an *in situ* hybridization screen for genes expressed in ways prefiguring the *y* gene expression pattern. We found that *wingless* (*wg*) expression precisely foreshadowed the six rows of black spots (Fig. 2b). Additionally, *decapentaplegic* (*dpp*) expression foreshadowed the dorsal and median pairs of spot rows (Fig. 2c), while *abdominal*-*A* (*abd-A*) expression correlated with the lateral pair of spot rows and the dorsal midline shade (Fig. 2d, e). *hedgehog* (*hh*) and *zerknullt* (*zen*) were additionally expressed along the dorsal midline of the abdomen (Fig. 2f, g). Thus, the activation of the *D. guttifera* color pattern appears to be induced in a modular fashion, which is in agreement with our observation that abdominal pigmentation patterns within the quinaria group are variations of the *D. guttifera* pattern ground plan (Fig. 3). This situation is reminiscent of the wing pattern ground plan in nymphalid butterflies^20,21^.

**Figure 2:**
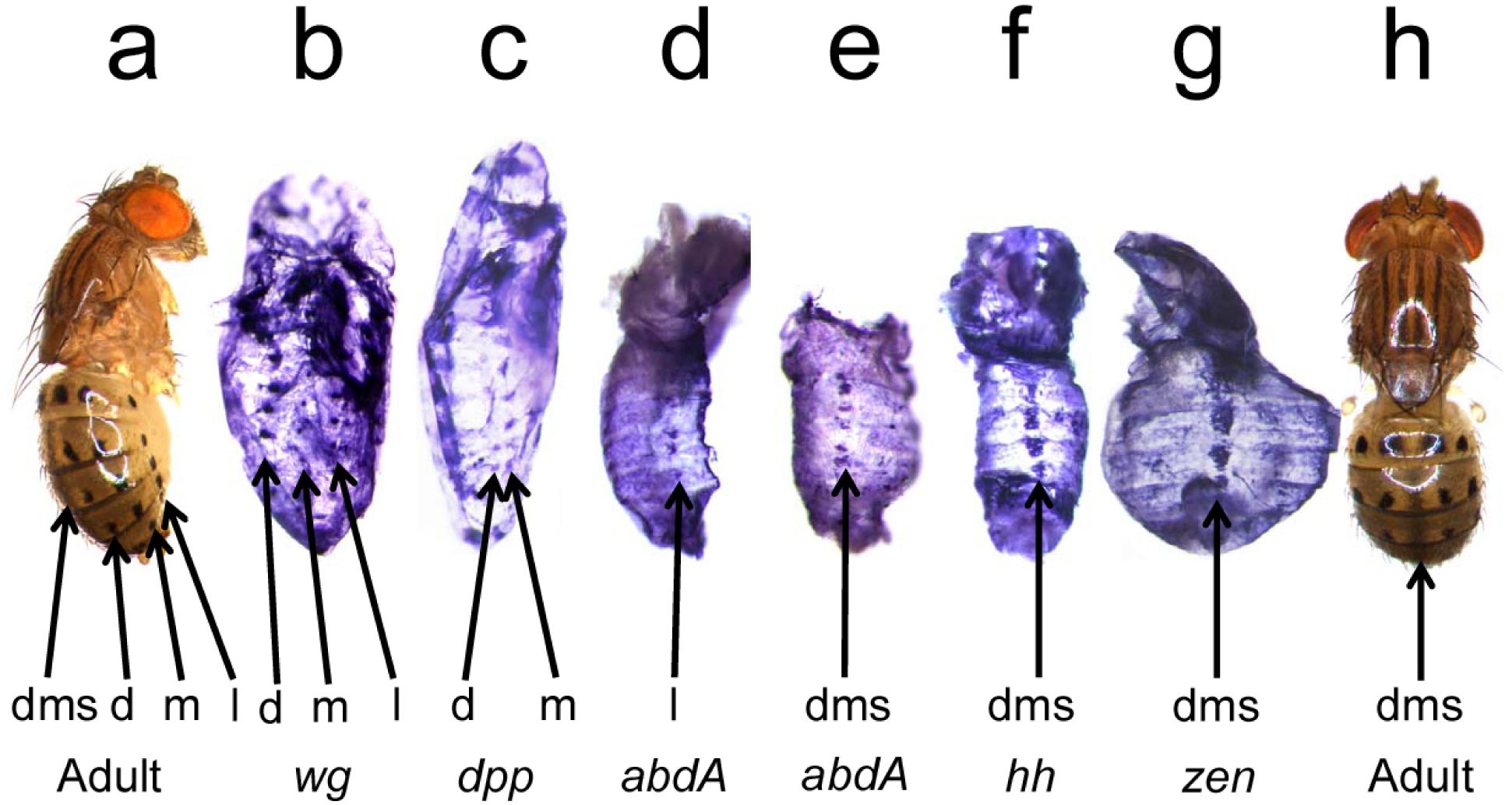
The mRNA expression patterns of five developmental genes foreshadow the *yellow* expression pattern. a, Adult, lateral view. b-g, *in situ* hybridizations in pupal abdomens. h, Adult, dorsal view. dms = dorsal midline shade, d = dorsal, m = median, l = lateral spot rows.

**Figure 3:**
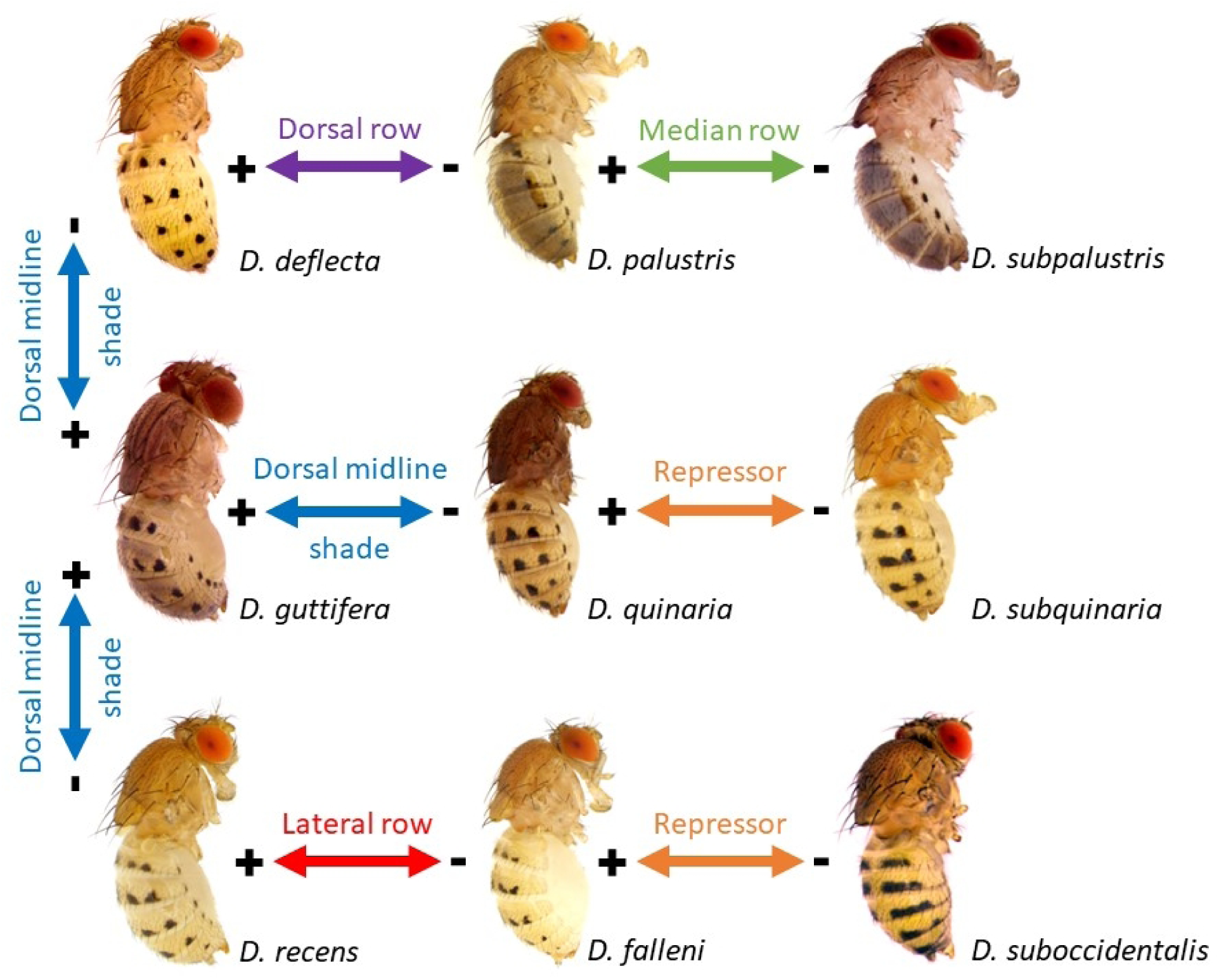
Deviations from the *D. guttifera* ground plan create the diversity of quinaria species’ abdominal color patterns. + = gain, - = loss of a pattern element. “Repressor” suggests stripes may be broken into spots by repressors of pigmentation and vice versa. The illustration does not imply any evolutionary direction; it solely illustrates the modularity of these complex patterns.

We hypothesized that the developmental candidate genes may activate the *y* gene through four CRMs, each controlling one sub-pattern to assemble the complete melanin pattern. We searched for these CRMs by transforming *D. guttifera* with *DsRed* reporter constructs containing non-coding fragments of the 42 kb *D. guttifera y* gene locus^12^ (Extended Data Fig. 1). Surprisingly, only one 953 bp fragment from the *y* intron, the *gut y spot* CRM, drove expression closely resembling all six spot rows on the developing abdomen (Fig. 4). To isolate possible sub-pattern-inducing CRMs, we subdivided the *gut y spot* CRM into 8 partially overlapping sub-fragments. Unexpectedly, the 636 bp left sub-fragment displayed horizontal stripe expression along the posterior edges of each abdominal segment, while the 570 bp right fragment was inactive (#1 & #2, Fig. 4.). Further dissection of this CRM revealed a 259 bp sub-fragment, which contained the minimal *gut y core stripe* CRM with some additional dorsal midline shade activity (#7, Fig. 4). These results suggest that the *D. guttifera* spots may have evolved from an ancestral stripe pattern that became partially repressed to isolate the spots. Currently, we cannot offer any direct evidence for specific candidate repressor genes. Neither the *in situ* hybridization experiments nor the bioinformatics analyses, using Jaspar, resulted in putative pigment stripe repressors.

**Fig. 4:**
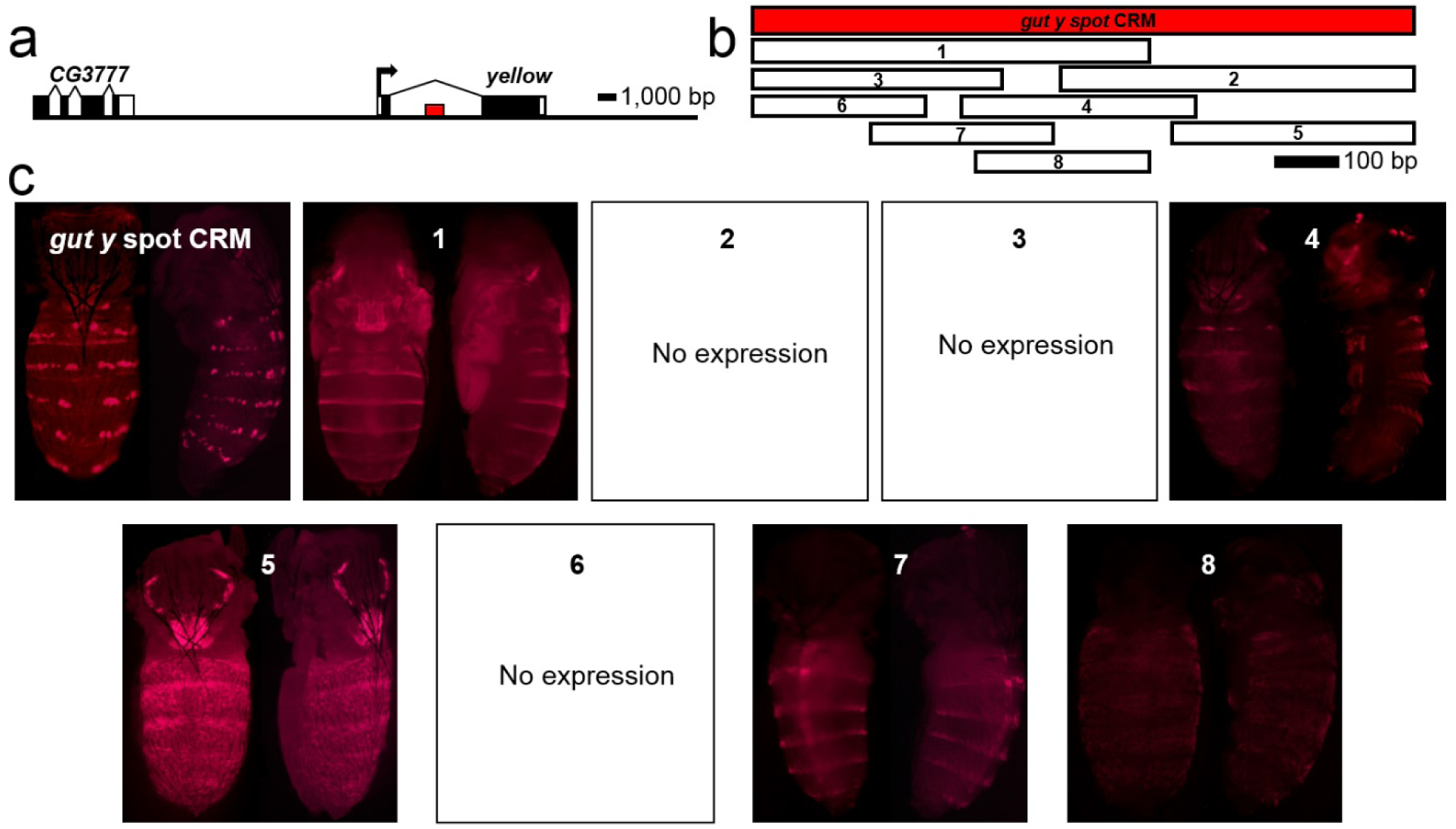
The *gut y spot* CRM is harbored within the *y* intron. a, The *y* gene locus. The red bar indicates the relative position of the *gut y spot* CRM. b, Sub-dividing the *gut y spot* CRM revealed horizontal stripes on each abdominal segment. The white bars (1-8) represent sub-fragments of the *gut y spot* CRM, and the corresponding pupal *DsRed* expression patterns in transgenic *D. guttifera* are shown.

Although we identified 24 Engrailed (En)-binding sites and 19 Homothorax (Hth)-binding sites in the *gut y spot* CRM (both are known repressors of pigmentation in *Drosophila*^15,22^), these sites were not enriched in the right half of the CRM, as we would have expected. However, our transcription factor binding site analysis of the *gut y spot* CRM sequence revealed putative transcription factor binding sites for most of the developmental genes that we identified as potential activators in our *in situ* hybridization screen, except for *hh*. This suggests that localized spot activation by these developmental factors contributes to the formation of the pattern.

**Extended Data Fig. 1:**
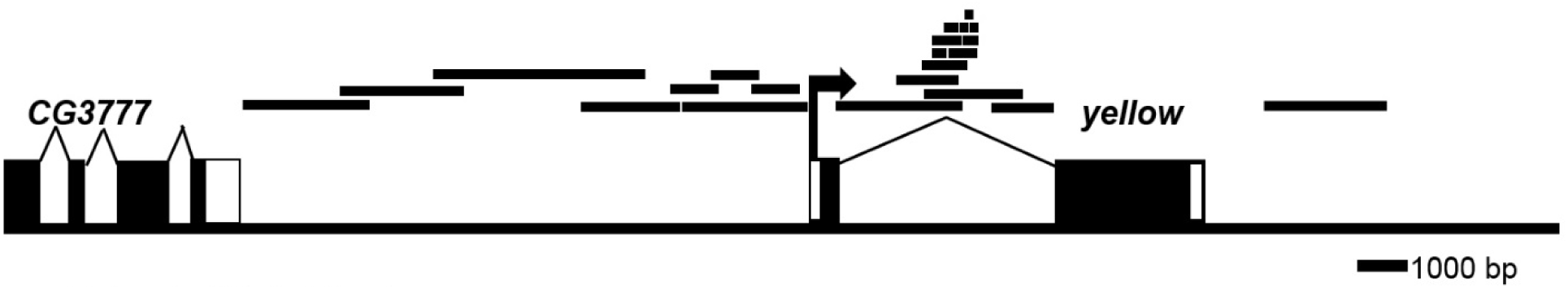
The *y* gene locus. The horizontal bars indicate the DNA fragments of the *D. guttifera y* gene that were tested in transgenic *D. guttifera* for regulatory activity.

Next, we asked whether the abdominal pigment spot pattern of a species closely related to *D. guttifera* shares a similar developmental basis. We thus performed *in situ* hybridization experiments in *Drosophila deflecta* (*D. deflecta)*. This species displays six longitudinal spot rows on its abdomen, but lacks the dorsal midline shade (Extended Data Fig. 2a, b). As in *D. guttifera, y* mRNA in *D. deflecta* pupal abdomens was expressed in six rows of spots, except along the dorsal midline (Extended Data Fig. 2c). Similarly, *wg* foreshadowed all six rows of spots, while *dpp* expression matched all but the lateral spot rows (Extended Data Fig. 3b, c). In contrast to the *D. guttifera* results, *abd-A, hh*, and *zen* were absent along the dorsal midline, which is in agreement with the lack of pigment in *D. deflecta* adults (Extended Data Fig. 3d, e, f, g). However, *abd-A* expression was not detectable where the lateral spot rows will form (Extended Data Fig. 3d), suggesting that these particular spots are controlled differently in *D. deflecta*. We next cloned the 938 bp orthologous *def y spot* CRM and transformed it into *D. guttifera*, using the *DsRed* reporter assay. The *def y spot* CRM drove faint dorsal spot row and stripe expression, especially along the dorsal spots (Extended Data Fig. 4). We further subdivided the *def y spot* CRM into 8 sub-fragments and identified a minimal *def y core stripe* CRM (288 bp) (#7, Extended Data Fig. 4). This sub-fragment drove a striped pattern, but without the dorsal midline shade activity seen in the *D. guttifera* minimal *gut y core stripe* CRM (#7, Fig. 4). We further transformed the *gut y spot* and *def y spot* CRMs including all sub-fragments into *D. melanogaster* to test if *D. melanogaster trans-*factors can bind to and activate these two quinaria group species’ spot CRMs. As a result, none of the reporter constructs showed any expression (data not shown). This suggests that the hypothetical ancestral stripe pattern of the quinaria group and the pigment stripes found on the *D. melanogaster* abdominal tergites^23^ have evolved independently by changes in *trans*. As the spot CRMs from *D. guttifera* and *D. deflecta* are not orthologous to any sequences within the *D. melanogaster y* locus, changes in *cis* have also contributed to the diversification of pigment patterns between *D. melanogaster* and the quinaria species group.

**Extended Data Fig. 2:**
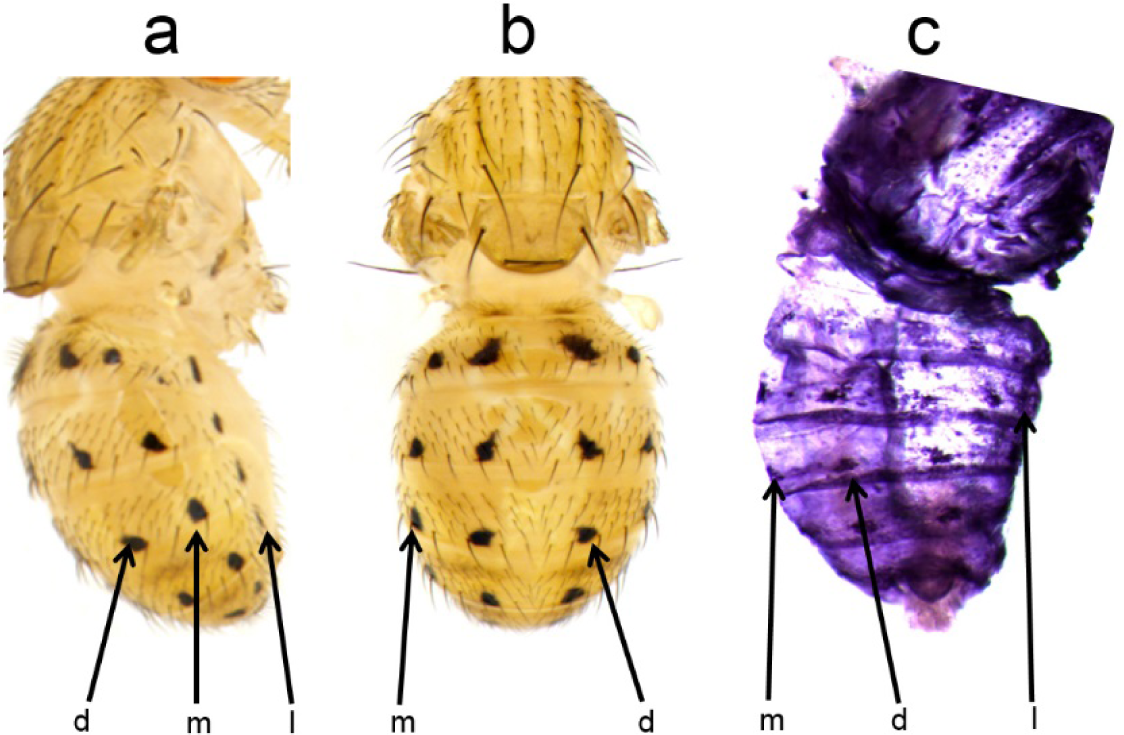
The *y* gene expression pattern in *D. deflecta* foreshadows the black spot pattern on the adult abdomen. a, Adult lateral view. b, Adult dorsal view. c, *y* mRNA in the pupal epidermis. d = dorsal, m = median, l = lateral.

**Extended Data Fig. 3:**
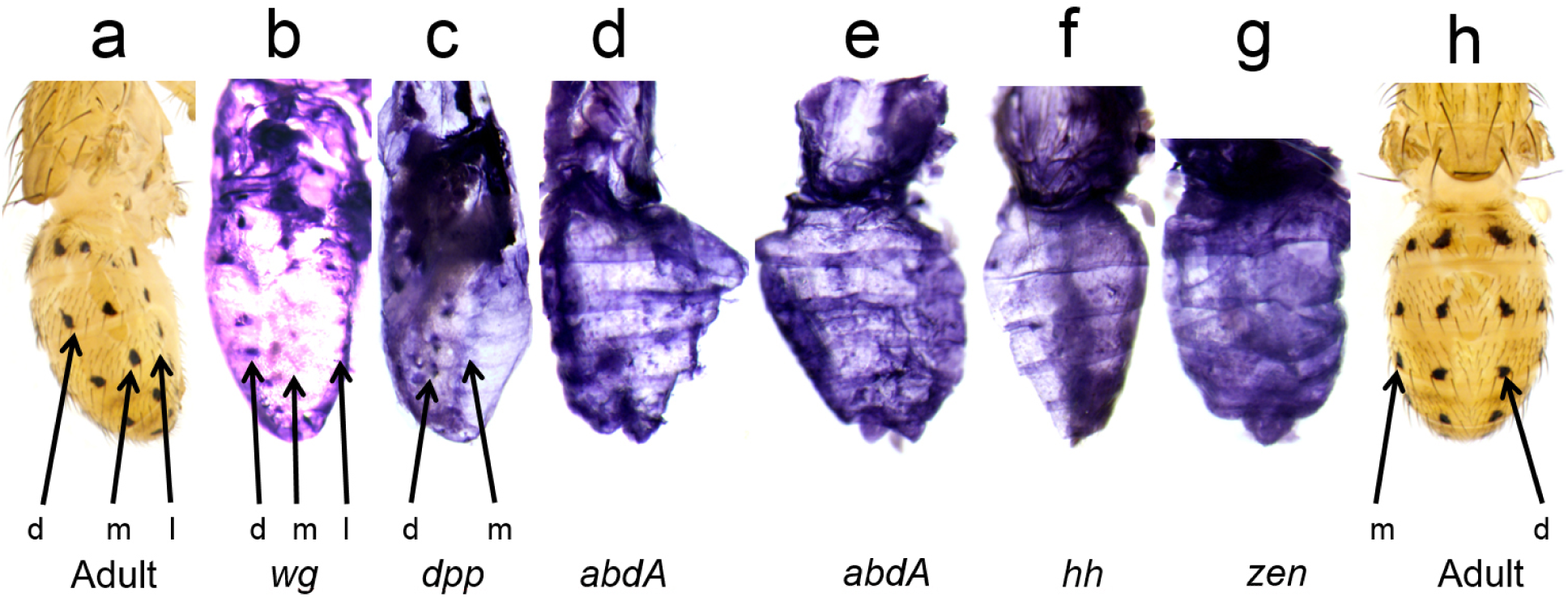
Developmental gene expression patterns in *D. deflecta* foreshadow distinct subsets of the adult abdominal color pattern. a, h, Adult, lateral view. b-g, Pupal *in situ* hybridizations. d = dorsal, m = median, l = lateral.

**Extended Data Fig. 4:**
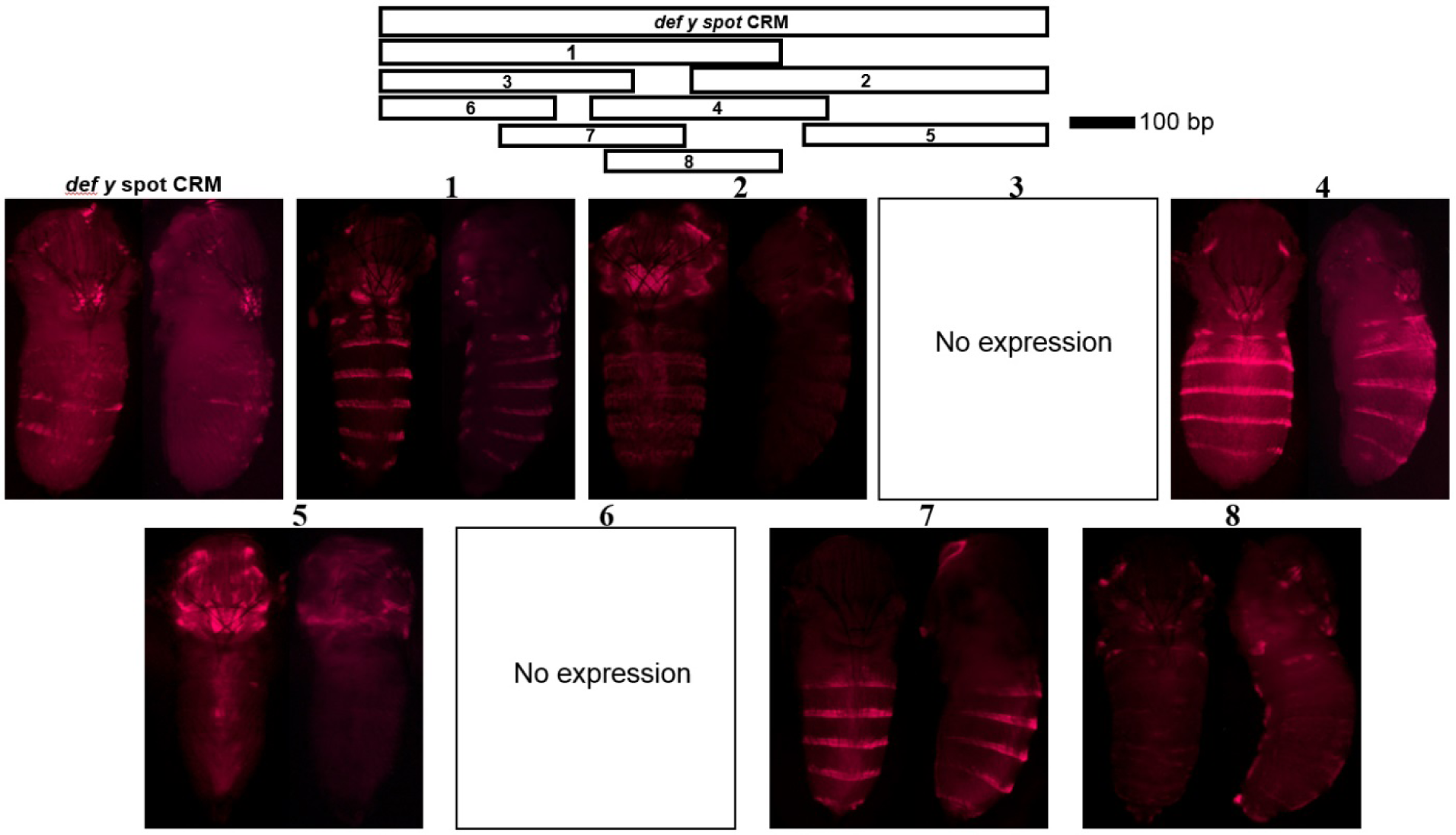
The orthologous *D. deflecta* region (*def y spot* CRM) analyzed in transgenic *D. guttifera.* The white bars (1-8) indicate sub-fragments of the *def y spot* CRM that were tested for reporter activity. The corresponding DsRed reporter expression patterns in developing pupae are shown.

In contrast to the *D. guttifera* wing spot pattern^12^, the abdominal pigment pattern develops in the absence of visible physical landmarks. *wg, dpp*, and *hh* are homologous to known proto-oncogenes in humans^24^, while *zen* and *abdA* are Hox genes. The abdominal color pattern of *D. guttifera* appears to be regulated by multiple developmental pathways consisting of activators and repressors acting in parallel, possibly targeting pigmentation genes other than *y* as well^18,19,25,26^. Further evidence for the repression of stripes can be seen in *Drosophila falleni*’s intraspecific pigment variation, another member of the quinaria species group (Extended Data Fig. 5). Our multi-pathway model fits well with the observation that the abdominal pattern variation presented by quinaria group members is largely due to modular derivations from the *D. guttifera* ground plan (Fig. 3). This scenario is reminiscent of the modularity found in butterfly wing patterns. Because insects use similar genes for color pattern development^21,27-30^, the quinaria group may serve as a valuable model to understand insect color pattern evolution. Future work should aim to manipulate the genes involved in pigmentation to test if they interact according to the reaction-diffusion model, as predicted by Alan Turing^31^.

**Extended Data Fig. 5:**
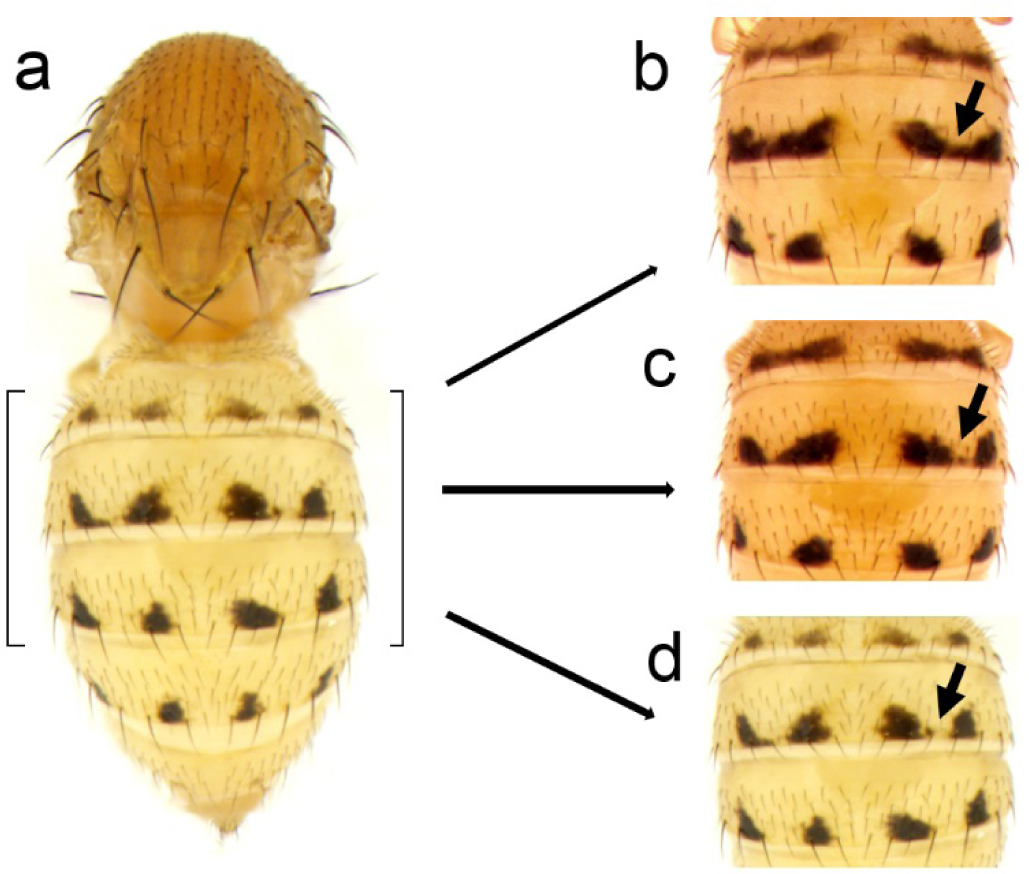
The abdominal pigment stripes of *D. falleni* break down into pigment spots. Intraspecific variation, as illustrated here, is very common in *D. falleni*. a, Adult abdomen. b-d, Adult abdominal pigment spots developing from stripe repression (arrows).

## Materials and Methods

### Molecular procedures

*In situ* hybridizations were carried out with species-specific RNA probes, as described in^12^, but with abdomens cut into halves and cleaned from the internal organs. At least three positive pupae were observed for each result shown. Additional images for verification purposes are provided in Extended Data Figs. 6-16. Immunohistochemistry for the Y protein in abdomens was performed according to^15^, with abdomens cut in half and cleaned with 1X PBS. *D. guttifera* CRMs were identified and tested in *D. guttifera* according to^12^ and in *D. melanogaster* as described in^23^. Transgenic experiments were performed as outlined in^32^. Pupal stages were identified according to^33^.

### *Drosophila* stocks

All fly stocks were a kind gift from the Sean B. Carroll Laboratory (University of Wisconsin - Madison) and were cultured at room temperature. We used the *D. melanogaster* fly strain VK00006 (cytogenic location 19E7), the *D. guttifera* stock no.15130–1971.10, and the *D. deflecta* stock no. 15130-2018.00 for gene expression and transgenic analyses.

### PCR primer sequences

We used the following primers to amplify the CRM sequences:

(iii) *gut y spot* CRM: Fwd: 5’-CAGCTGCGGTTGAGTACGAC-3’and Rvs: 5’-GCCAACTCGACGGGAATTC-3’. Restriction sites: KpnI and SacII.

(iv) *def y spot* CRM: Fwd: 5’-CAGCTGCTGCGGTTCAGTAG-3’ and Rvs: 5’-GCTAGACACACGTTGGTTTGCT-3’. Restriction sites: KpnI and SacII.

(v) *gut y spot* CRM sub-fragment #1: Fwd: 5’-CAGCTGCGGTTGAGTACGAC-3’ and Rvs: 5’-ACTGAATCTGATTTCGGCTCG-3’. Restriction sites: KpnI and SacII.

(vi) *gut y spot* CRM sub-fragment #2: Fwd: 5’-AGTTAATCGCCAGTCAATAATGGC-3’ and Rvs: 5’-GAATTCCCGTCGAGTTGGC-3’. Restriction sites: KpnI and SacII.

(vii) *gut y spot* CRM sub-fragment #3: Fwd: 5’-CAGCTGCGGTTGAGTACGAC-3’ and Rvs: 5’-GCCATTATTGACTGGCGATTAAC-3’. Restriction sites: KpnI and SacII.

(viii) *gut y spot* CRM sub-fragment #4: Fwd: 5’-AAATGAAGCTCAGTGAGCCGC-3’ and Rvs: 5’-ACTGAATCTGATTTCGGCTCG-3’. Restriction sites: KpnI and SacII.

(ix) *gut y spot* CRM sub-fragment #5: Fwd: 5’-AGCATCTGAAACTTAAACGCCG-3’ and Rvs: 5’-GAATTCCCGTCGAGTTGGC-3’. Restriction sites: KpnI and SacII.

(x) *gut y spot* CRM sub-fragment #6: Fwd: 5’-CAGCTGCGGTTGAGTACGAC-3’ and Rvs: 5’-CAGCGATATTAATTTTTTATTCAATGG-3’. Restriction sites: KpnI and SacII.

(xi) *gut y spot* CRM sub-fragment #7(*gut y core stripe* CRM): Fwd: 5’-AAATGAAGCTCAGTGAGCCGC-3’ and Rvs: 5’-GCGATTTGTTTGTCAAGTCAAC-3’. Restriction sites: KpnI and SacII.

(xii) *gut y spot* CRM sub-fragment #8: Fwd: 5’-AAATGAAGCTCAGTGAGCCGC-3’ and Rvs: 5’-GTTGACTTGACAAACAAATCGC-3’. Restriction sites: KpnI and SacII.

(xiii) *def y spot* CRM sub-fragment #1: Fwd: 5’-CAGCTGCTGCGGTTCAGTAG-3’ and Rvs: 5’-ATTGTCGCAGCTGCCTAACG-3’. Restriction sites: KpnI and SacII.

(xiv) *def y spot* CRM sub-fragment #2: Fwd: 5’-AACGAAGCTCACTGAGCTGC-3’ and Rvs: 5’-AGCAAACCAACGTGTGTCTAGC-3’. Restriction sites: KpnI and SacII.

(xv) *def y spot* CRM sub-fragment #3: Fwd: 5’-CAGCTGCTGCGGTTCAGTAG-3’ and Rvs: 5’-GTTAAAAGCAGCCAGTTGGCC-3’. Restriction sites: KpnI and SacII.

(xvi) *def y spot* CRM sub-fragment #4: Fwd: 5’-CAAAGAATCGAATTCGGAGACAG-3’ and Rvs: 5’-ATTGTCGCAGCTGCCTAACG-3’. Restriction sites: KpnI and SacII. (Clone name: *def y* 1.1C2)

(xvii) *def y spot* CRM sub-fragment #5: Fwd: 5’-GAATGAGATTCGTTAGGCAGC-3’ and Rvs: 5’-AGCAAACCAACGTGTGTCTAGC-3’. Restriction sites: KpnI and SacII.

(xviii) *def y spot* CRM sub-fragment #6: Fwd: 5’-CAGCTGCTGCGGTTCAGTAG-3’ and Rvs: 5’-TTCAACGGATATTCGTTCAATTTC-3’. Restriction sites: KpnI and SacII.

(xix) *def y spot* CRM sub-fragment #7 (*def y core stripe* CRM): Fwd: 5’-CAAAGAATCGAATTCGGAGACAG-3’ and Rvs: 5’-GTCAGGCAATGTAAATGTTGTCG-3’. Restriction sites: KpnI and SacII.

(xx) *def y spot* CRM sub-fragment #8: Fwd: 5’-AACGAAGCTCACTGAGCTGC-3’ and Rvs: 5’-ATTGTCGCAGCTGCCTAACG-3’. Restriction sites: KpnI and SacII.

These forward and reverse primer sequences do not include restriction sites.

**Extended Data Fig. 6:**
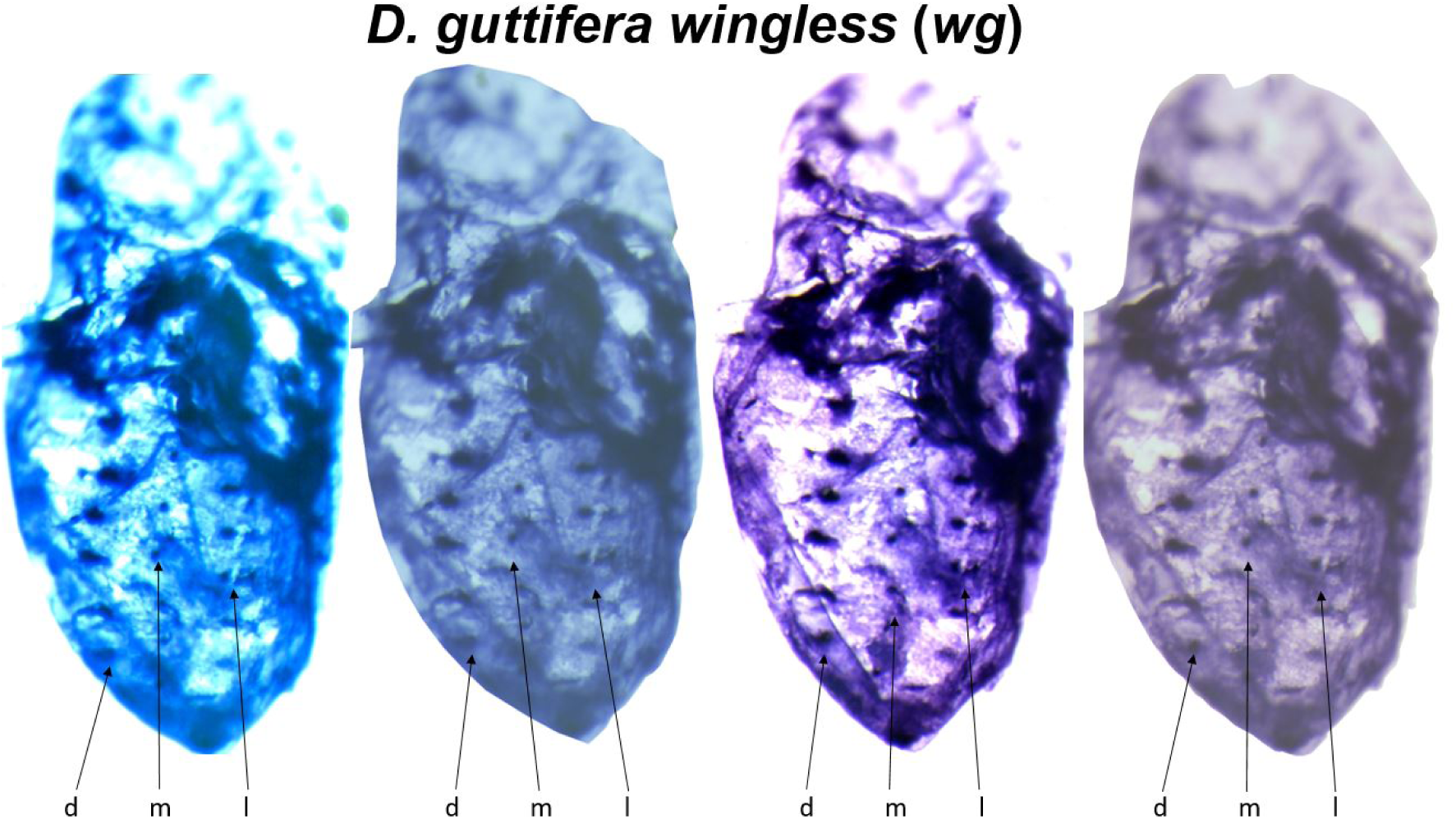
One *D. guttifera* pupa stained with a *wg* probe. d = dorsal, m = median, l = lateral row of spots. Different image manipulations shown.

**Extended Data Fig. 7:**
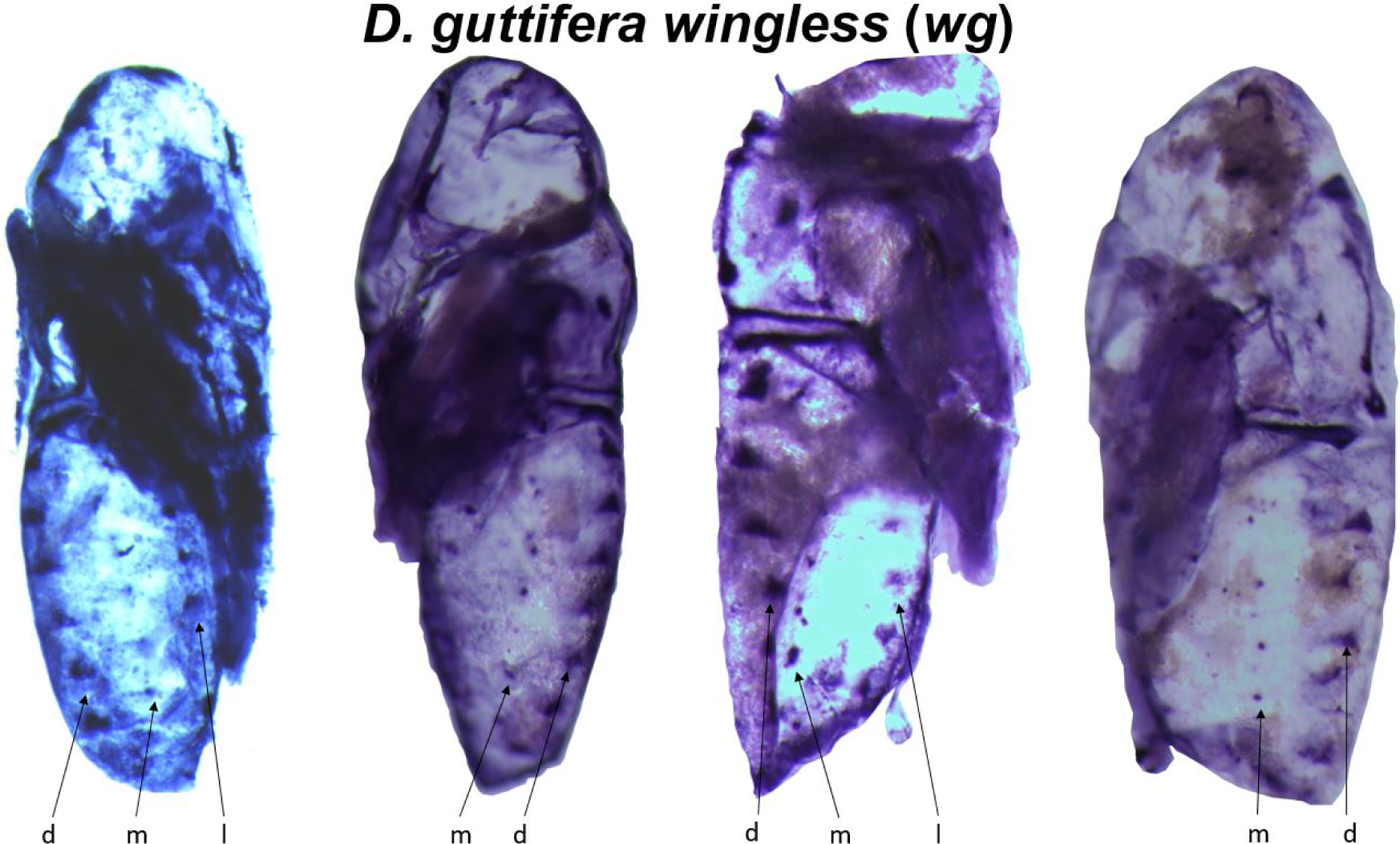
*D. guttifera* pupae stained with a *wg* probe. d = dorsal, m = median, l = lateral row of spots.

**Extended Data Fig. 8:**
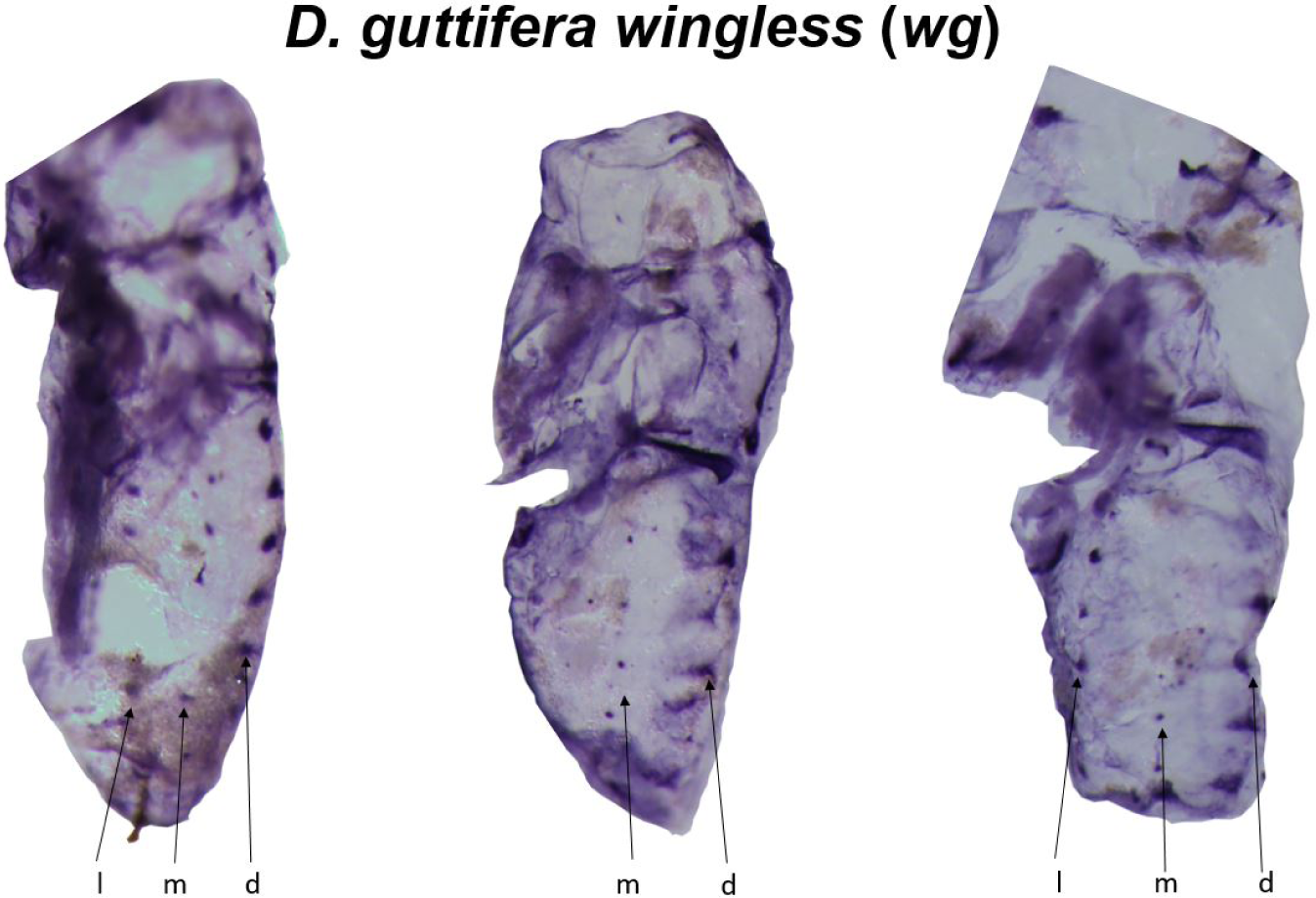
*D. guttifera* pupae stained with a *wg* probe. d = dorsal, m = median, l = lateral row of spots.

**Extended Data Fig. 9:**
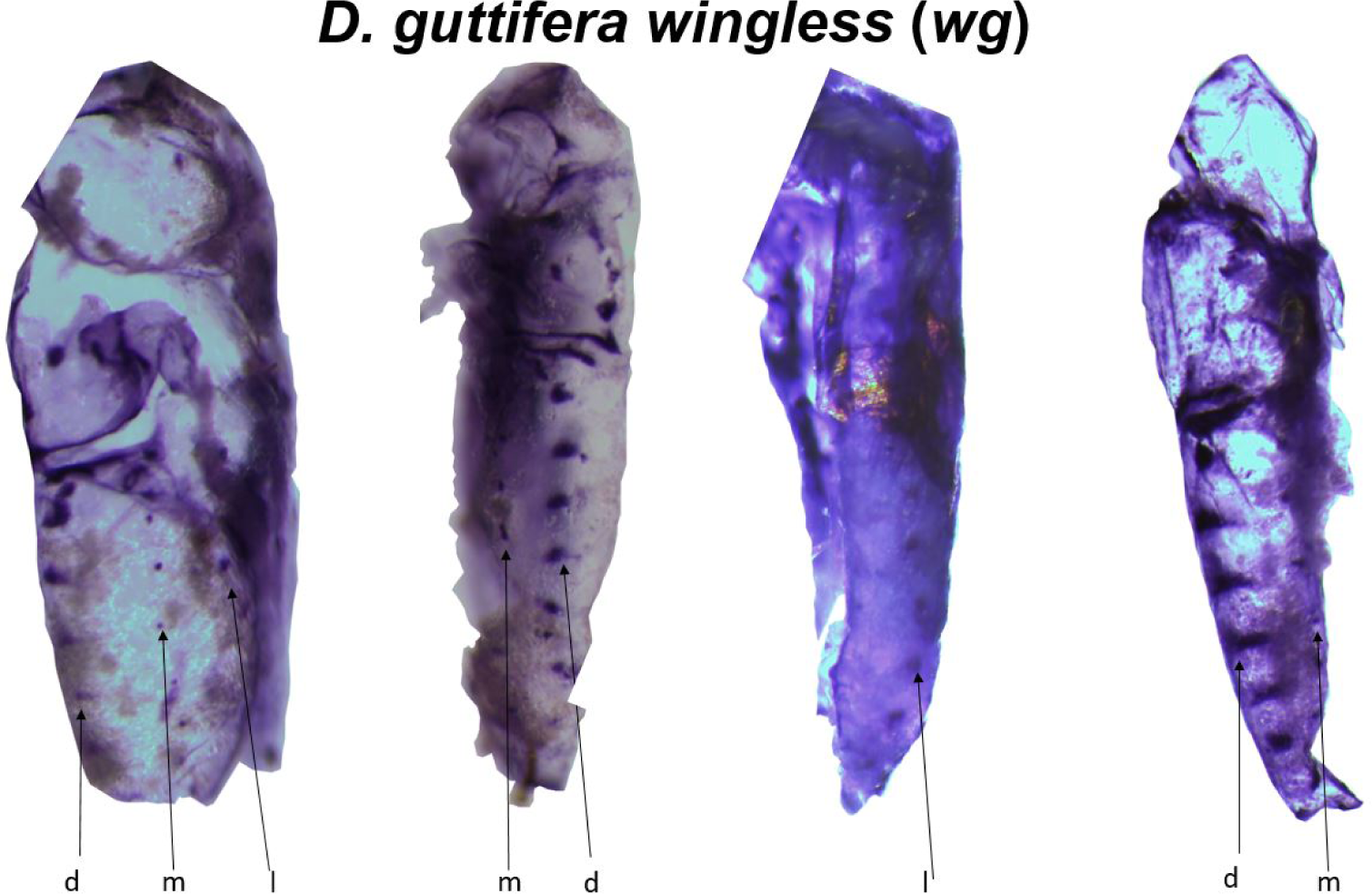
*D. guttifera* pupae stained with a *wg* probe. d = dorsal, m = median, l = lateral row of spots.

**Extended Data Fig. 10:**
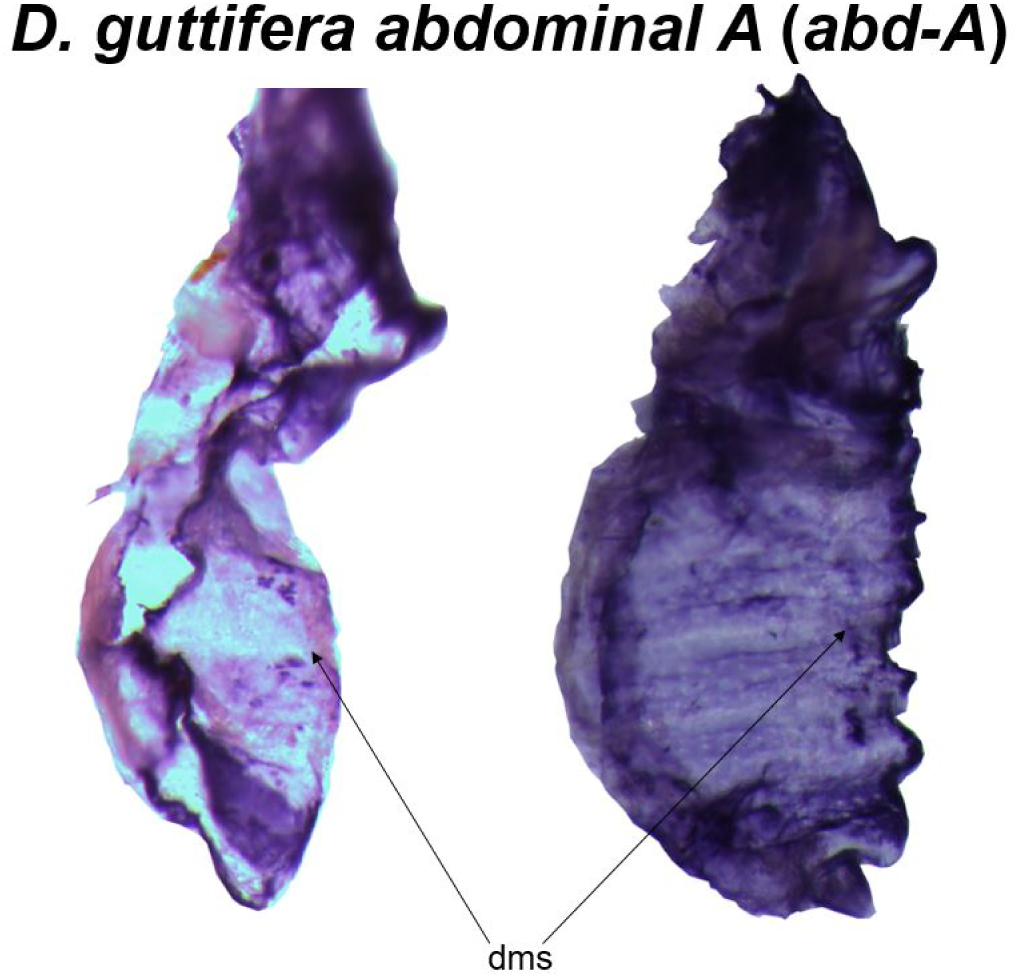
*D. guttifera* pupae stained with an *abd-A* probe. dms = dorsal midline shade.

**Extended Data Fig. 11:**
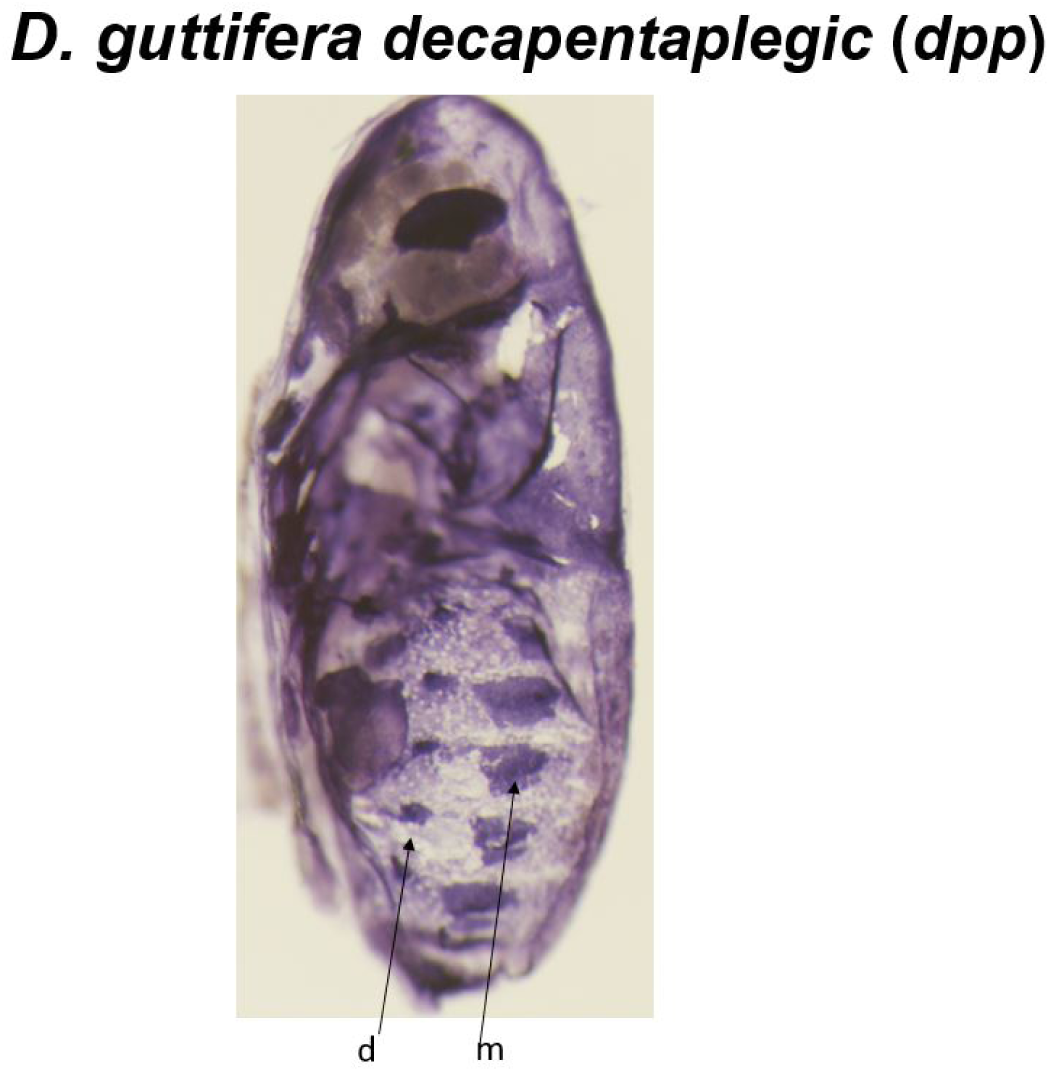
*D. guttifera* pupa stained with a *dpp* probe. d = dorsal, m = median row of spots.

**Extended Data Fig. 12:**
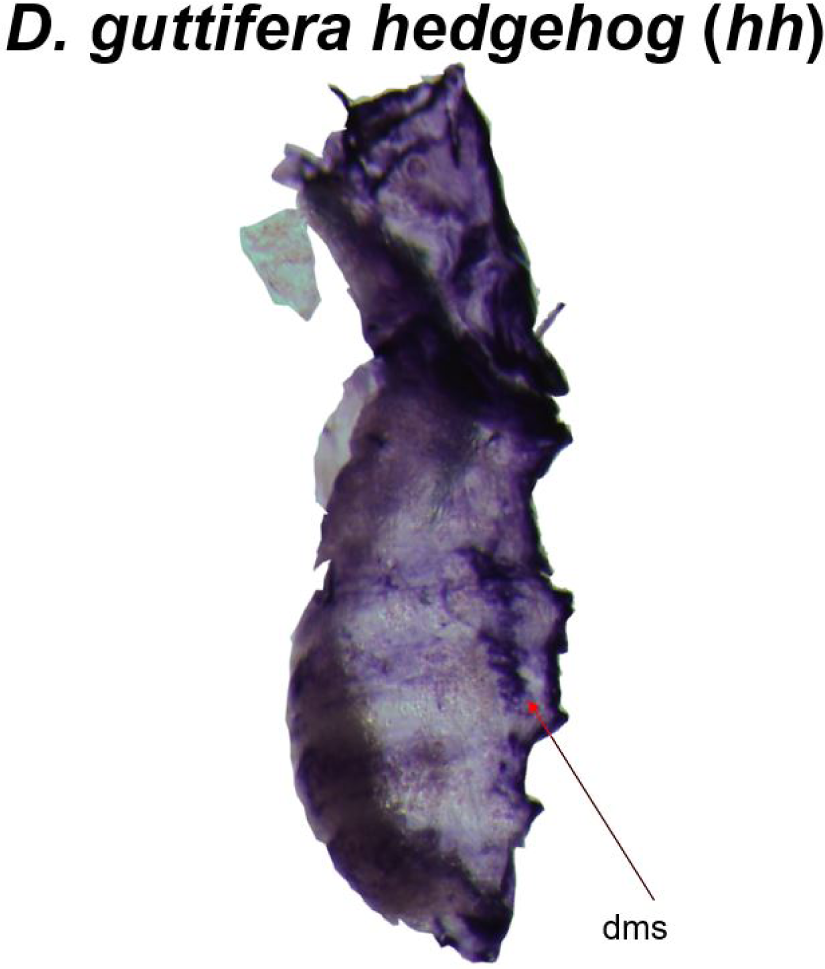
*D. guttifera* pupa stained with a *hh* probe. dms = dorsal midline shade.

**Extended Data Fig. 13:**
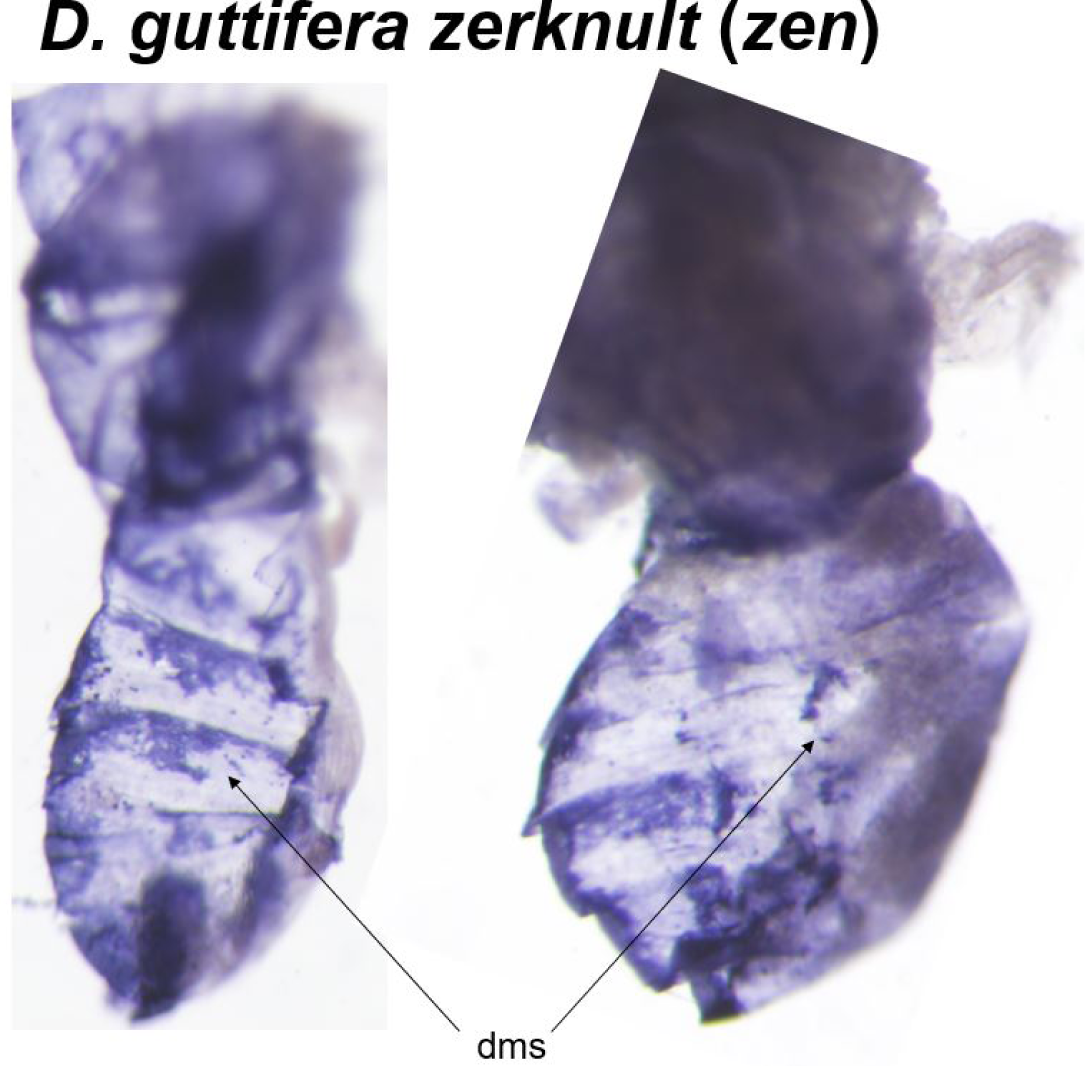
*D. guttifera* pupae stained with a *zen* probe. dms = dorsal midline shade.

**Extended Data Fig. 14:**
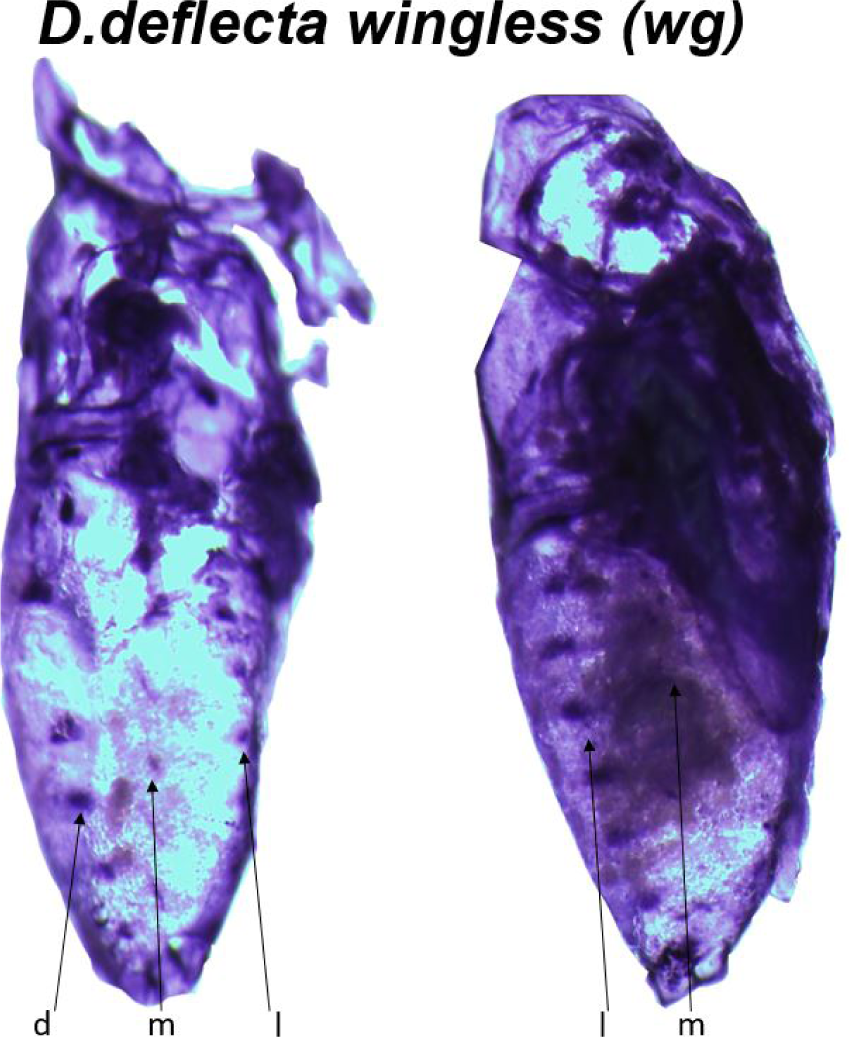
*D. deflecta* pupae stained with a *wg* probe. d = dorsal, m = median, l = lateral row of spots.

**Extended Data Fig. 15:**
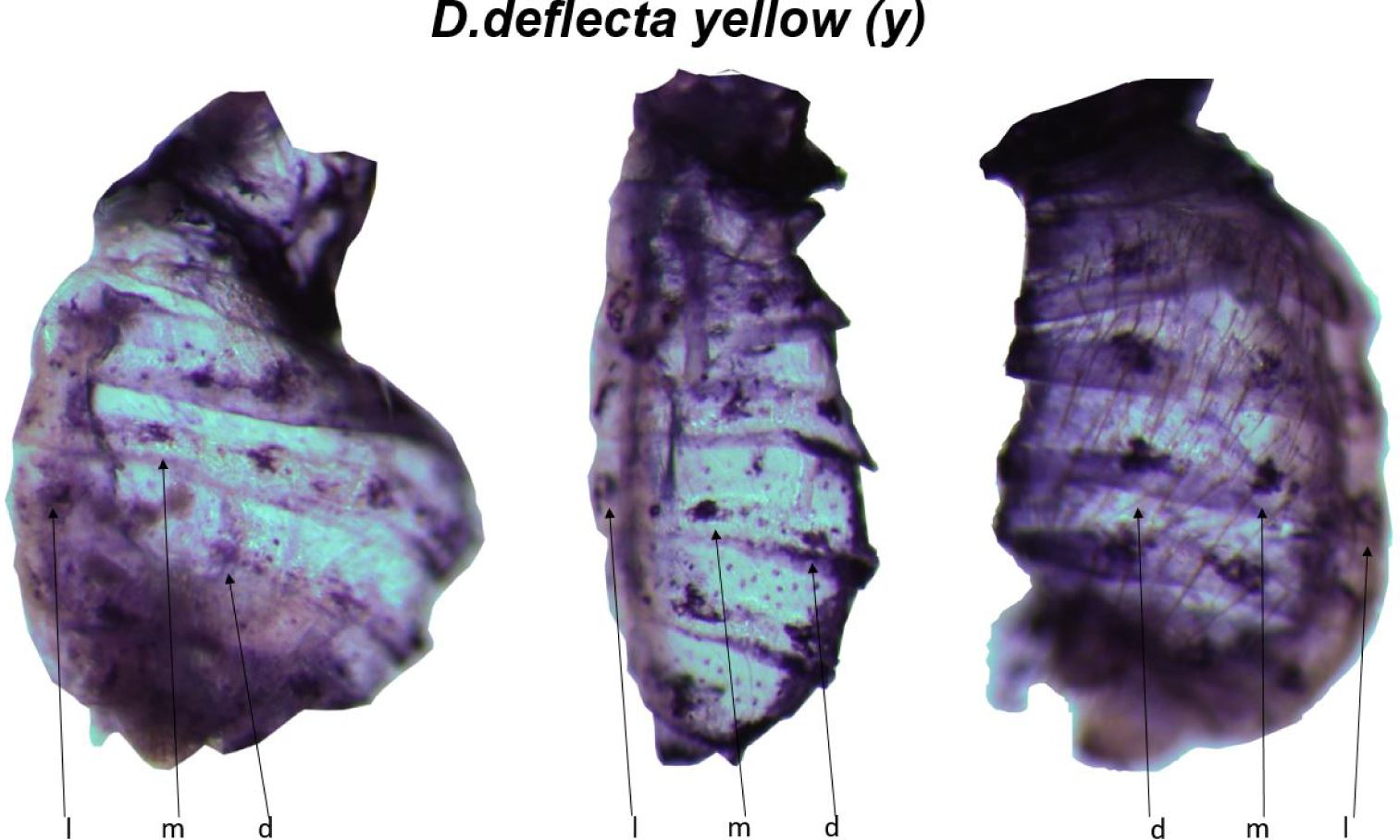
*D. deflecta* pupae stained with a *y* probe. d = dorsal, m = median, l = lateral row of spots.

**Extended Data Fig. 16:**
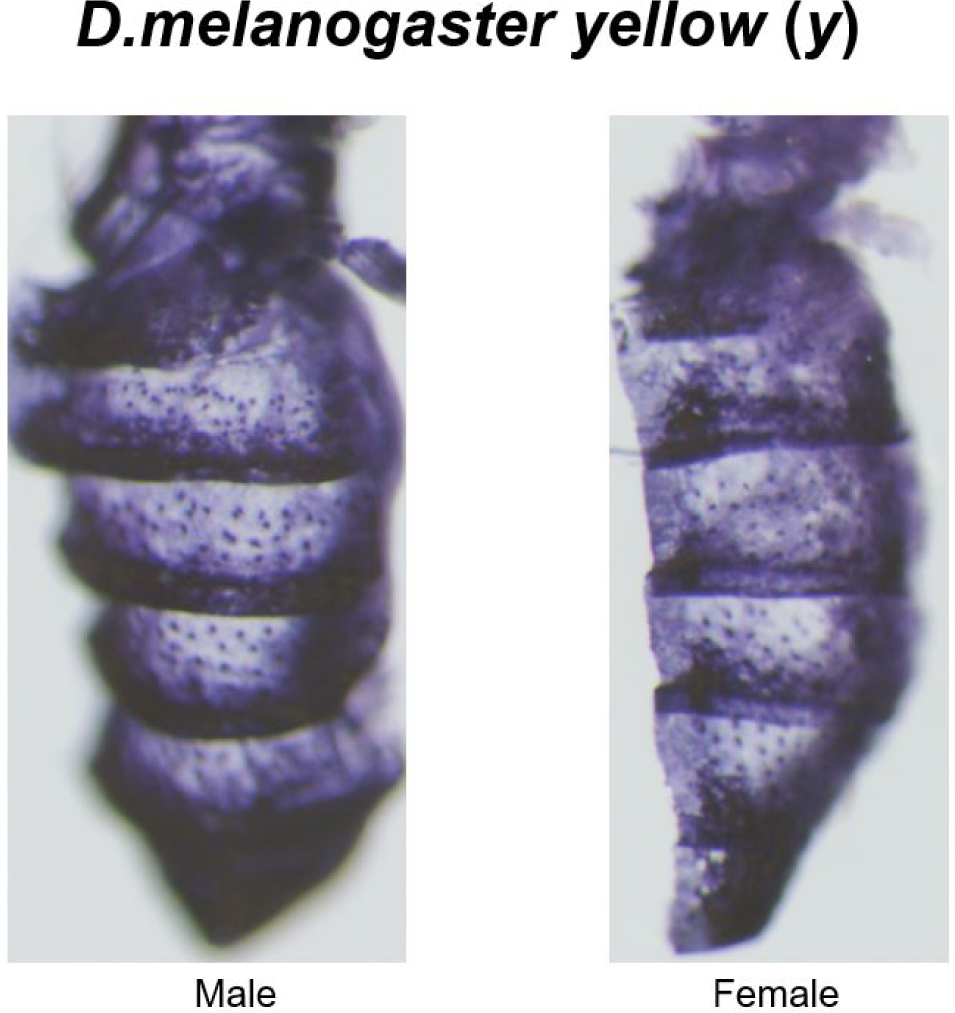
*D. melanogaster* pupae stained with a *y* probe.

## Acknowledgements

We thank Ryan Bensen, Abigail Meisel, Bridgette Rebbeck, and Jason Hu for technical assistance. This research was funded by NIH grant #1R15GM107801-01A1 to T.W.

## Author Contributions

K.K.B.R., M.S. and T.W. conceived and designed the experiments; K.K.B.R, P.M.E.N., T.E.S, E.B., P.P.K, A.McQ., E.M., A.A., A.N. and T.W. performed the experiments; K.K.B.R., S.M. and T.W. analyzed the data; T.W. obtained funding; K.K.B.R and T.W. wrote the paper; K.K.B.R, S.M., P.M.E.N., T.E.S., E.B., P.P.K., A.McQ., E.M., A.A., A.N. and T.W. edited the paper.

## References

1 True, J. R. & Carroll, S. B. Gene co-option in physiological and morphological evolution. Annu Rev Cell Dev Bi. 18, 53–80 (2002).

2 Carroll, S. B. Endless forms: the evolution of gene regulation and morphological diversity. Cell 101, 577–580 (2000).

3 McGregor, A. P. et al. Morphological evolution through multiple *cis*-regulatory mutations at a single gene. Nature 448, 587-U586, doi:10.1038/nature05988 (2007).

4 Shapiro, M. D. et al. Genetic and developmental basis of evolutionary pelvic reduction in threespine sticklebacks. Nature 428, 717–723 (2004).

5 Rubinstein, M. & de Souza, F. S. Evolution of transcriptional enhancers and animal diversity. Philos Trans R Soc Lond B Biol Sci. 368, 20130017, doi:10.1098/rstb.2013.0017 (2013).

6 McKay, D. J., Estella, C. & Mann, R. S. The origins of the *Drosophila* leg revealed by the *cis*-regulatory architecture of the Distalless gene. Development 136, 61-71, doi:10.1242/dev.029975 (2009).

7 Werner, T., Steenwinkel, T., Jaenike, J. (2018) Drosophilids of the Midwest and Northeast. (Version 2) J. Robert Van Pelt and John and Ruanne Opie Library, Michigan Technological University. Houghton, Michigan. ISBN: 978-1-7326524-0-8 (E-Book, 345 pages) https://digitalcommons.mtu.edu/oabooks/1/

8 Bray, M. J., Werner, T. & Dyer, K. A. Two genomic regions together cause dark abdominal pigmentation in *Drosophila tenebrosa*. Heredity 112, 454-462, doi:10.1038/hdy.2013.124 (2013).

9 True, J. R. Insect melanism: the molecules matter. Trends Ecol Evol. 18, 640-647, doi:DOI 10.1016/j.tree.2003.09.006 (2003).

10 Wittkopp, P. J., Vaccaro, K. & Carroll, S. B. Evolution of *yellow* gene regulation and pigmentation in *Drosophila*. Curr Biol. 12, 1547–1556 (2002).

11 Riedel, F., Vorkel, D. & Eaton, S. Megalin-dependent *yellow* endocytosis restricts melanization in the *Drosophila* cuticle. Development 138, 149-158, doi:10.1242/dev.056309 (2011).

12 Werner, T., Koshikawa, S., Williams, T. M. & Carroll, S. B. Generation of a novel wing colour pattern by the Wingless morphogen. Nature 464, 1143-1148, doi:nature08896[pii] 10.1038/nature08896 (2010).

13 Gibert, J. M., Mouchel-Vielh, E. & Peronnet, F. Modulation of *yellow* expression contributes to thermal plasticity of female abdominal pigmentation in *Drosophila melanogaster*. Sci Rep. 7, 43370, doi:10.1038/srep43370 (2017).

14 Jeong, S., Rokas, A. & Carroll, S. B. Regulation of body pigmentation by the Abdominal-B Hox protein and its gain and loss in *Drosophila* evolution. Cell 125, 1387-1399, doi:S0092-8674(06)00760-4 [pii]10.1016/j.cell.2006.04.043 (2006).

15 Gompel, N., Prud’homme, B., Wittkopp, P. J., Kassner, V. A. & Carroll, S. B. Chance caught on the wing: *cis*-regulatory evolution and the origin of pigment patterns in *Drosophila*. Nature 433, 481–487 (2005).

16 Prud’homme, B. et al. Repeated morphological evolution through *cis*-regulatory changes in a pleiotropic gene. Nature 440, 1050–1053 (2006).

17 Arnoult, L. et al. Emergence and diversification of fly pigmentation through evolution of a gene regulatory module. Science 339, 1423-1426, doi:10.1126/science.1233749339/6126/1423 [pii] (2013).

18 Ordway, A. J., Hancuch, K. N., Johnson, W., Wiliams, T. M. & Rebeiz, M. The expansion of body coloration involves coordinated evolution in *cis* and *trans* within the pigmentation regulatory network of *Drosophila prostipennis*. Dev Biol. 392, 431-440, doi:10.1016/j.ydbio.2014.05.023 (2014).

19 Camino, E. M. et al. The Evolutionary Origination and Diversification of a Dimorphic Gene Regulatory Network through Parallel Innovations in *cis* and *trans*. Plos Genet. 11, doi:ARTN e1005136 10.1371/journal.pgen.1005136 (2015).

20 Nijhout, H. F. & Wray, G. A. Homologies in the Color Patterns of the Genus *Heliconius* (Lepidoptera, Nymphalidae). Biol J Linn Soc. 33, 345-365, doi:DOI 10.1111/j.1095-8312.1988.tb00449.x (1988).

21 Sekimura, T. & Nijhout, H. F. Diversity and Evolution of Butterfly Wing Patterns: An Integrative Approach. (Springer, 2017).

22 Rogers, W. A. et al. A survey of the *trans*-regulatory landscape for *Drosophila melanogaster* abdominal pigmentation. Dev Biol 385, 417-432, doi:10.1016/j.ydbio.2013.11.013 (2014).

23 Williams, T. M. et al. The regulation and evolution of a genetic switch controlling sexually dimorphic traits in *Drosophila*. Cell 134, 610–623 (2008).

24 Geissler, K. & Zach, O. Pathways involved in *Drosophila* and human cancer development: the Notch, Hedgehog, Wingless, Runt, and Trithorax pathway. Ann Hematol. 91, 645-669, doi:10.1007/s00277-012-1435-0 (2012).

25 Miyagi, R., Akiyama, N., Osada, N. & Takahashi, A. Complex patterns of *cis*-regulatory polymorphisms in *ebony* underlie standing pigmentation variation in *Drosophila melanogaster*. Mol Ecol. 24, 5829-5841, doi:10.1111/mec.13432 (2015).

26 Dion, W. A., Shittu, S. O., Steenwinkel, T. E., Raja, K. K. B., Kokate, P. P., & Werner, T. From simplicity to complexity: The gain or loss of spot rows underlies the morphological diversity of three *Drosophila* species. bioRxiv. doi: https://doi.org/10.1101/2020.04.03.024778 (2020).

27 Martin, A. & Reed, R. D. Wnt signaling underlies evolution and development of the butterfly wing pattern symmetry systems. Dev Biol. 395, 367-378, doi:10.1016/j.ydbio.2014.08.031 (2014).

28 Ozsu, N., Chan, Q. Y., Chen, B., Das Gupta, M. & Monteiro, A. Wingless is a positive regulator of eyespot color patterns in *Bicyclus anynana* butterflies. Dev Biol. 429, 177-185, doi:10.1016/j.ydbio.2017.06.030 (2017).

29 Futahashi, R., Banno, Y. & Fujiwara, H. Caterpillar color patterns are determined by a two-phase melanin gene prepatterning process: new evidence from *tan* and *laccase2*. Evol Dev. 12, 157-167, doi: EDE401 [pii] 10.1111/j.1525-142X.2010.00401.x (2010).

30 Miyazaki, S. et al. Sexually dimorphic body color is regulated by sex-specific Expression of *yellow* gene in ponerine ant, *Diacamma* sp. Plos One 9, doi:ARTN e92875 10.1371/journal.pone.0092875 (2014).

31 Turing, A. M. The chemical basis of morphogenesis. Phil Trans R Soc B. 237, 37–72 (1952).

32 Shittu, M., Steenwinkel, T., Koshikawa, S. & Werner, T. The making of transgenic *Drosophila guttifera*. Preprints. 2020040120 (doi:10.20944/preprints202004.0120.v1) (2020).

33 Fukutomi, Y., Matsumoto, K., Agata, K., Funayama, N. & Koshikawa, S. Pupal development and pigmentation process of a polka-dotted fruit fly, *Drosophila guttifera* (Insecta, Diptera). Dev Genes Evol 227, 171–180 (2017).

